# Widespread microbial mercury methylation genes in the global ocean

**DOI:** 10.1101/648329

**Authors:** Emilie Villar, Lea Cabrol, Lars-Eric Heimbürger-Boavida

**Affiliations:** Aix Marseille Université, Univ Toulon, CNRS, IRD, Mediterranean Institute of Oceanography (MIO) UM 110, 13288, Marseille, France; Sorbonne Université, Université Pierre et Marie Curie - Paris 6, CNRS, UMR 7144 (AD2M), Station Biologique de Roscoff, Place Georges Teissier, CS90074, Roscoff, 29688 France

**Author notes:** Corresponding Author: Léa Cabrol. Both authors contributed equally to this work (Lea Cabrol, Lars-Eric Heimbürger-Boavida).

## Abstract

Methylmercury is a neurotoxin that bioaccumulates from seawater to high concentrations in marine fish, putting human and ecosystem health at risk. High methylmercury levels have been found in the oxic subsurface waters of all oceans, yet only anaerobic microorganisms have been identified so far as efficient methylmercury producers in anoxic environments. The microaerophilic nitrite oxidizing bacteria *Nitrospina* has been previously suggested as a possible mercury methylator in Antarctic sea ice. However, the microorganisms processing inorganic mercury into methylmercury in oxic seawater remain unknown. Here we show metagenomic evidence from open ocean for widespread microbial methylmercury production in oxic subsurface waters. We find high abundances of the key mercury methylating genes *hgcAB* across all oceans corresponding to taxonomic relatives of known mercury methylators from Deltaproteobacteria, Firmicutes and Chloroflexi. Our results identify *Nitrospina* as the predominant and widespread key player for methylmercury production in the oxic subsurface waters of the global ocean.

## Introduction

Human activities release 2500 tons of inorganic mercury (Hg) every year and have added 55 000 tons of Hg to the global ocean since the industrial revolution ^1^. Humans are exposed to Hg in the form of methylmercury (MeHg), mainly through the consumption of marine fish. The Minamata Convention (www.mercuryconvention.org) aims to protect human health from the adverse effects of Hg *via* the reduction of anthropogenic, inorganic Hg emissions. To understand the efficacy and time-scales of lowered Hg emissions to reduce fish MeHg levels, we must fully understand the origin of marine MeHg. Microorganisms play a central role in Hg transformations. We must identify the Hg methylating microbes and the factors controlling their distribution in order to better constrain MeHg production in the global ocean.

As the only cultured microbes known to produce MeHg to date are anaerobic, research focused for many years on a MeHg source in anoxic marine sediments ^2–5^. However, several lines of independent evidence speak in favour of *in situ* MeHg production in oxic seawater as the main source of fish MeHg. Recent large scale oceanographic expeditions found subsurface MeHg maxima in every ocean basin ^4,6^. The proportion of MeHg to inorganic Hg throughout the oxic seawater column is higher than those found in anoxic sediments. Laboratory experiments show that Hg methylation can occur in anaerobic microniches that occur within sinking particles in oxic waters ^7^. Bianchi et al. ^8^ provide compelling evidence that anaerobic microbes thrive in anoxic microenvironments of sinking particulate organic matter. Independently, incubation experiments with isotopically labelled Hg spikes show significant *in situ* Hg methylation in oxic seawater ^9^. Additional evidence stems from Hg stable isotope signatures of marine fish, that can only be explained if 60-80% of the MeHg is produced in open ocean subsurface waters ^10^. Lastly, a pioneering study found a compound specific δ^13^C signature of fish tissue MeHg similar to algal δ^13^C, suggesting that MeHg is produced in the open ocean water column ^11^.

A major breakthrough has been made with the discovery of two key genes, *hgcA* and *hgcB*, that control Hg methylation in model anaerobic Hg-methylators^5^. The presence of the *hgcAB* operon predicts Hg methylation capacity in diverse microorganisms ^2^. A screening of publicly available microbial metagenomes found the *hgcAB* genes in nearly all anaerobic environments, but the study only rarely detected the genes in pelagic marine water column metagenomes in the open ocean ^12^. In antarctic sea ice a marine microaerophilic nitrite oxidizing bacterium belonging to the *Nitrospina* genus has been recently identified as a potential Hg methylator with HgcA-like proteins ^13^. We aim to resolve the paradox between the wealth of geochemical evidence for *in situ* MeHg production and the absence of known anaerobic Hg methylators in the open ocean. Metagenomic data from 243 *Tara* Oceans samples from 68 open ocean locations covering most ocean basins was analysed to generate an ocean microbial reference gene catalog^14^. We screened the *Tara* Oceans metagenomes for the presence of the key *hgcA* methylating gene and provide compelling evidence on the potential key players producing MeHg in the open ocean.

## Results and Discussion

### Identification of HgcAB homologs in the ocean gene catalog

Ten *hgcA* and 5 *hgcB* homolog genes were identified in the Ocean Microbial Reference Gene Catalog ^14^ (OM-RGC), 6 scaftigs presenting simultaneously *hgcA* and *hgcB* (Fig. 1, Supplementary Table 1). Alignment of HgcA sequences revealed 7 sequences with the conserved NVWCAA motif ^5^ and one sequence with the modified NIWCAA motif on the ‘cap helix’ region. Mutation experiments previously showed that the structure of the putative ‘cap helix’ region harbouring Cys93 is crucial for methylation function ^15^. Two HgcA sequences were truncated (OM-RGC.v1.019516181, OM-RGC.v1.015822836), preventing inspection of their conserved motif, but they could be unequivocally assigned to HgcA sequences based on their phylogenetic placement and high similarity with known HgcA sequences (Fig. 2). The 5 HgcB sequences presented the conserved motif ECGAC ^5^ (Supplementary Table 1).

**Figure 1.**
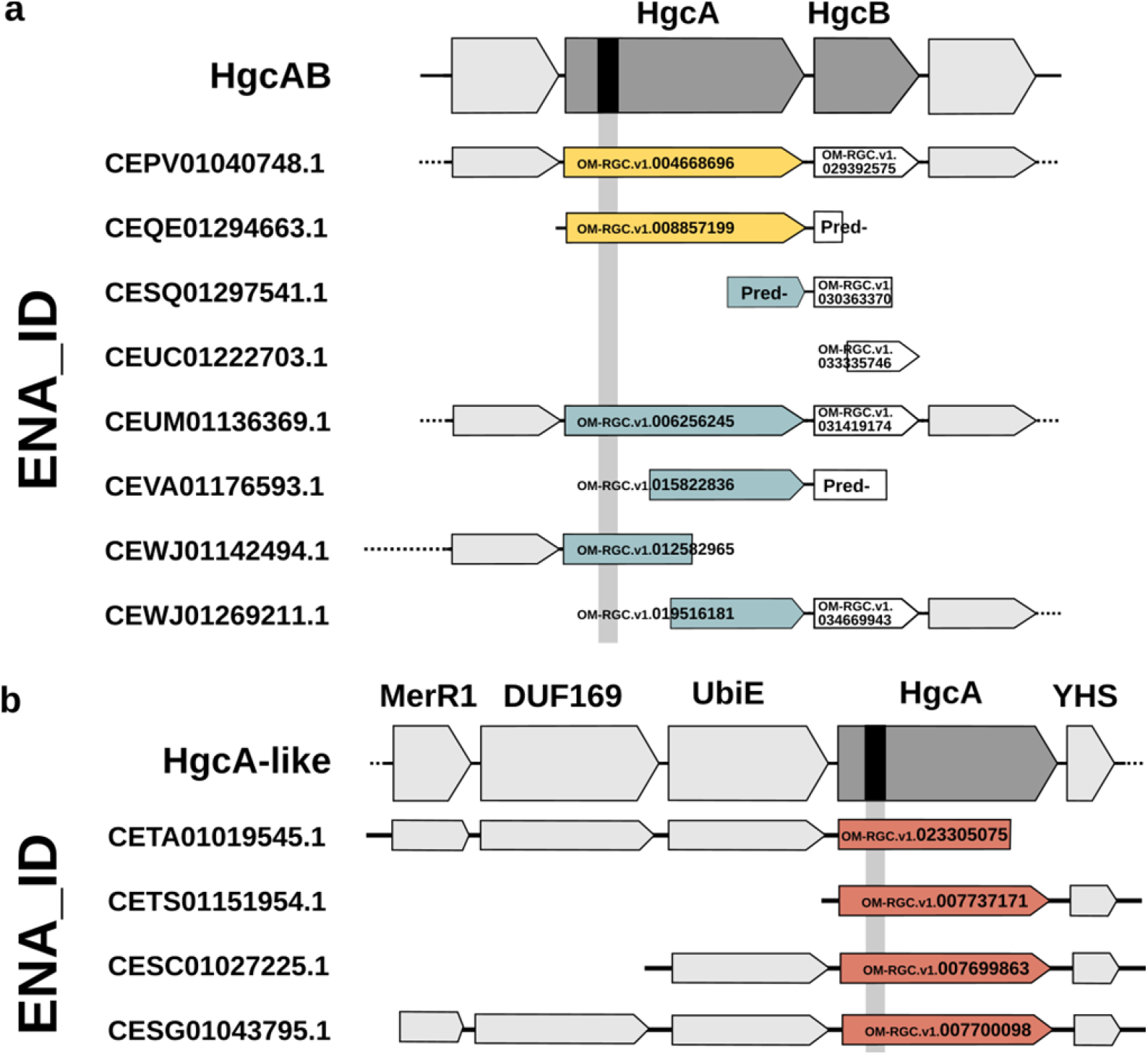
The genomic context of the HgcA orthologs. **a**, HgcAB operon. **b**, HgcA-like proteins. The 12 retrieved scaftigs are identified by their ENA_ID on the left of the figure. The solid lines represent the extent of the scaftig sequence and the dashed lines indicate that the scaftig sequence is longer than the represented part. The location of the conserved motif is indicated on the HgcA box by a black bar. When present in *Tara* Oceans samples, the corresponding gene identifier is indicated on the bar for HgcA and HgcB, or indicated as (Pred-) if the gene was incomplete and the protein sequence was partially predicted. The colour of the HgcA boxes refers to the biogeographical clustering as defined in Fig.3 (Cluster 1 in blue, Cluster 2 in yellow, Cluster 3 in red). For Cluster 3 sequences (assigned to Nitrospina), the genomic context was enlarged to show the closest sequences (MerR1: mercuric resistance operon regulatory protein, UbiE: Ubiquinone/menaquinone biosynthesis C-methyltransferase, DUF169: Hypothetical protein with DUF 169 motif, YHS: Hypothetical protein with YHS domain).

**Figure 2.**
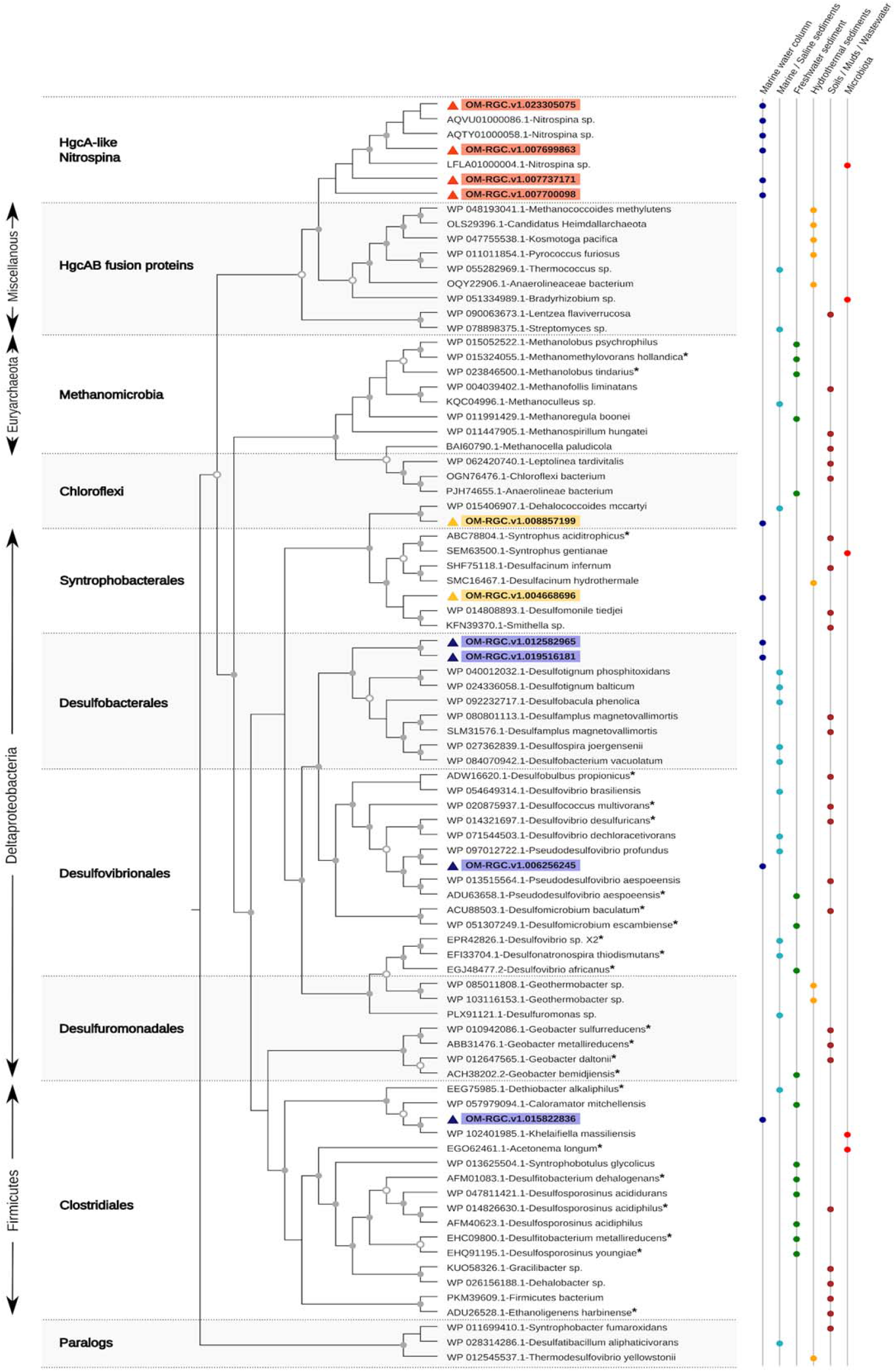
Phylogenetic tree of HgcA homolog sequences found in the *Tara* Oceans assemblies. Maximum likelihood phylogenies were inferred using PhyML Best AIC Tree with the best model of sequence evolution Blosum62+I+G+F. Branch support was calculated using the non-parametric Shimodaira-Hasegawa-like approximate likelihood ratio test. The triangle colour refers to the biogeographical clustering of the HgcA sequences retrieved from *Tara* Oceans assemblies, as defined in Fig.3 (Cluster 1 in blue, Cluster 2 in yellow, Cluster 3 in red). The tree was rooted using 3 paralogues from confirmed non-Hg methylating bacteria. Sequences from experimentally confirmed mercury methylators were indicated with an asterisk. Support values using 1,000 resamples are shown when >50 and coloured squares indicate the isolation source.

### HgcA sequences found in the Tara Oceans assemblies covered nearly all known Hg methylators

Phylogenetic placement of the 10 sequences found in the *Tara* Oceans assemblies covered nearly all known Hg methylators (Fig. 2). Four sequences (OM-RGC.v1.007700098, OM-RGC.v1.007737171, OM-RGC.v1.023305075, OM-RGC.v1.007699863) were closely related to the HgcA-like proteins described by Gionfriddo et al. ^13^ for *Nitrospina* sp. The Nitrospinae phylum has been described as a distinct phylogenetic group of lithoautotrophic nitrite oxidizing bacteria exclusively found in marine environments ^16^, particularly abundant in oxygen-deficient zones ^17^.

The remaining 6 HgcA sequences were distributed between Deltaproteobacteria, Firmicutes and Chloroflexi phyla. Within Deltaproteobacteria, three orders were represented, namely *Desulfovibrionales, Desulfobacterales* and *Syntrophobacterales.* OM-RGC.v1.006256245 was most closely related to HgcA from *Pseudodesulfovibrio profundus*, a strictly anaerobic piezophilic sulfate-reducer bacteria (SRB) previously isolated from marine sediment ^18^. It belongs to the Desulfovibrionales order, which contains several members with confirmed Hg-methylating capacity, such as the model species *Desulfovibrio desulfuricans* with exceptionally high Hg-methylation rates, isolated from estuarine sediments ^19^. OM-RGC.v1.019516181 and OM-RGC.v1.012582965 belonged to *Desulfobacterales*, a well-known order of SRB containing efficient Hg-methylators such as *Desulfobulbus propionicus* and *Desulfococcus multivorans*. Finally, OM-RGC.v1.004668696 belonged to Syntrophobacterales. The closest relative of OM-RGC.v1.004668696 with strong confirmed methylation potential was the non-SRB obligate syntroph *Syntrophus aciditrophicus* ^2^. Syntrophic bacteria are important Hg-methylators in low-sulfate ecosystems ^20,21^, where they degrade OM in association with H_2_-consuming microorganisms such as sulfate-reducers, iron-reducers and methanogens.

Within Firmicutes, OM-RGC.v1.015822836 was tightly related to HgcA from recently isolated human gut bacteria *Khelaifiella* in the Clostridiales order ^22^. Their closest relative with confirmed methylation potential is the non-SRB *Dethiobacter alkaliphilus*, with low to moderate Hg-methylation capacity ^2^.

OM-RGC.v1.008857199 was related to Chloroflexi, a phylum for which several *hgcAB*-carriers have been identified, but for which experimental confirmation of Hg methylation capacity is still needed. This sequence clusters tightly with HgcA from *Dehalococcoides mccartyi*, which has been reported as a potential methylator, albeit in minor abundance, in freshwater marshes ^20^. These two sequences are separated from other Chloroflexi HgcA sequences and more closely related to HgcA sequences from Syntrophobacterales, showing that the taxonomy and the HgcA-phylogeny are not always congruent. The phylogenetically irregular distribution of *hgcA* can be an indication of horizontal gene transfers (HGT) and/or gene deletions in response to stress, suggesting the prevalent influence of environment on Hg-methylation ability ^23^.

Among the 10 HgcA sequences found in the gene ocean catalogue, none was affiliated to methanogenic *Archaea*. Even if the co-existence of methanogens and sulfate-reducers has been evidenced in marine sediments ^24^, sulfate reduction usually outcompetes methanogenesis in seawater under non-limiting sulfate concentrations ^25^. Our results thus show that Hg-methylators in the ocean span a large taxonomic diversity, not limited to sulfate-reducing bacteria.

### Biogeography distinguishes three groups of putative marine Hg methylators

Once clearly identified and phylogenetically assigned, the biogeographic distribution patterns of *hgcA* was evaluated. The 10 HgcA sequences were identified in 77 samples out of the 243 available *Tara* Oceans metagenomes and were clearly distributed in three clusters according to their abundance patterns (Fig. 3). The biogeographic clustering was consistent with the HgcA-phylogeny.

**Figure 3.**
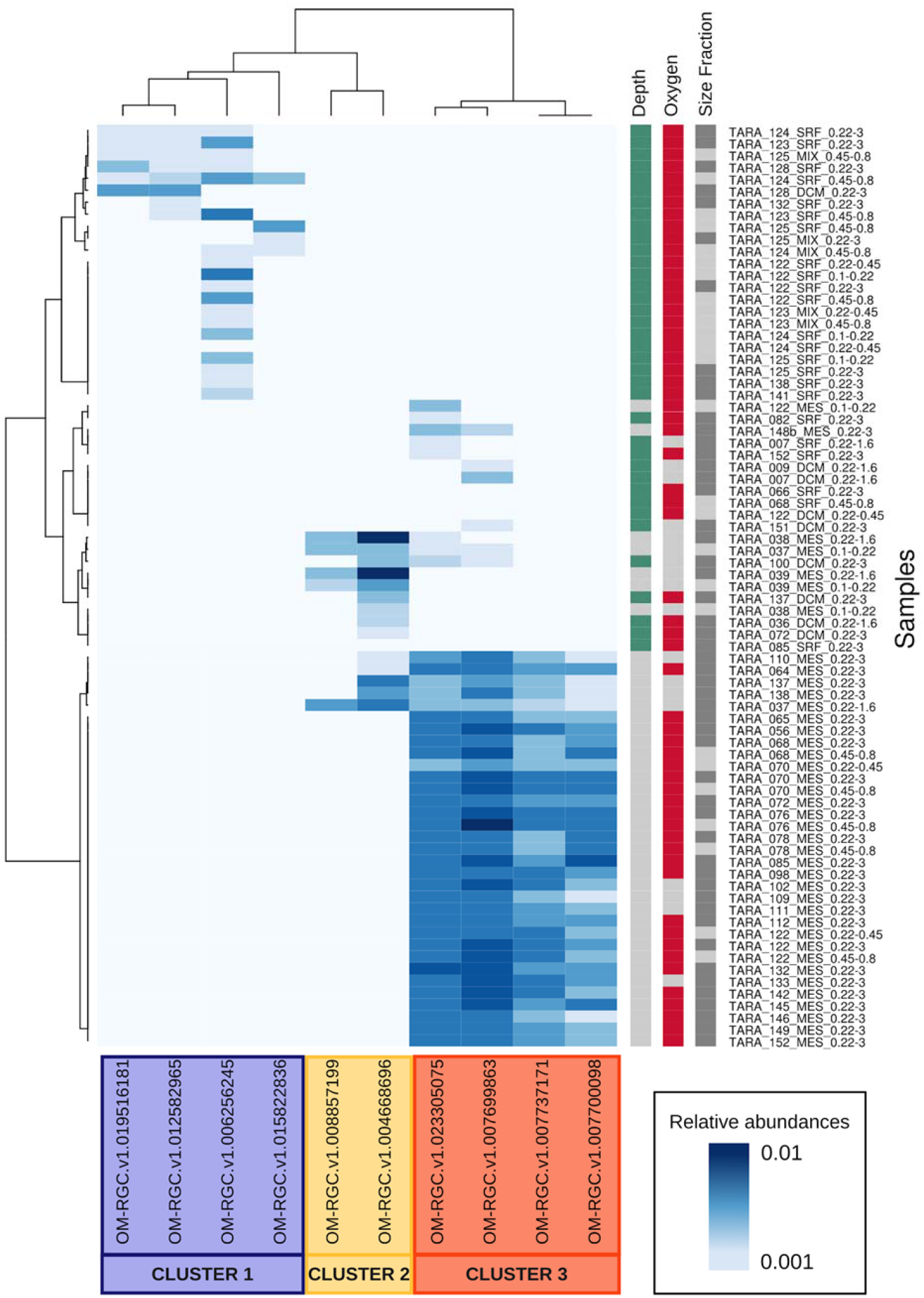
Distribution of HgcA in *Tara* Oceans samples. HgcA relative abundance (from 0.01 to 0.1) is indicated by the white-blue gradient. The hierarchical clustering highlighted three gene clusters with high abundances in specific samples with marked environmental features, as suggested by colored squares. Surface samples were collected in the upper layer (< 120 m-depth, in green) while subsurface were collected below 120 m-depth (in grey). Seawater was considered as oxic when O_2_ > 10 µM (in red) and suboxic when O_2_ < 10 µM (in grey). Larger size fraction samples are in dark grey (0.22-3 µm) and smaller size fractions samples (<0.8 µm) are in light grey.

Cluster 1 gathered Desulfobacterales, Clostridiales and Desulfovibrionales HgcA sequences, exclusively present in 23 oxic surface waters (< 120 m-depth, > 10 µM_O2_). Highest abundances were found in the photic zone of the Pacific Ocean, especially in the area surrounding the Marquesas Islands. This region is characterized by extensive plankton blooms triggered by a physico-chemical phenomenon called Island Mass Effect related to iron fertilization. In this Cluster, the HgcA sequence OM-RGC.v1.006256245 related to the *Desulfovibrionales* order (containing most of the experimentally confirmed Hg-methylators) was the most frequent and abundant in the 23 oxic samples.

The phylogenetic placement of the two sequences grouped in Cluster 2 is poorly supported. The most abundant sequence was related to *Smithella* and *Desulfomonile tiedjei* (Syntrophobacterales) while the other one was close to Chloroflexi (Fig. 2, Supplementary Table 2). HgcA sequences from Cluster 2 were identified in 15 surface and subsurface samples, mostly in suboxic waters: sequences found in samples with oxygen concentration below 10 µM accounted for 98% of total Cluster 2 abundances (Supplementary Figure 2). The highest abundances of Cluster 2 HgcA sequences were found in the subsurface waters of the northern stations within the Arabian Sea Oxygen Minimum Zone (Stations TARA_036 to TARA_039), under the influence of a previous major bloom event, where high particle concentrations and strong anaerobic microbial respiration have been reported ^26^. Cluster 2 sequences were also found in lower abundance in the shallow anoxic zone of the Pacific North Equatorial Counter Current (Stations TARA_137 and TARA_138, see methods).

The most abundant HgcA-like proteins were grouped in Cluster 3 and were exclusively assigned to *Nitrospina*. These *Nitrospina* HgcA-like proteins were found in 47 samples, widespread across all sampled ocean basins. They were almost exclusively found in subsurface water (> 120 m-depth) and were more frequent in the oxic waters (> 10 µM_O2_). Subsurface oxic waters accounted for 84% of total *Nitrospina-*HgcA abundance (Supplementary Figure 2). Their highest relative abundance was found in the South Atlantic and the South Pacific Oceans (Fig. 4, Supplementary Table 2). *Nitrospina* HgcA abundance was positively correlated to nitrate concentration (R 0.54, P < 0.005), which is consistent with *Nitrospina’s* role as nitrate producer through nitrite oxidation, and with the well-known nitrate enrichment with depth in the ocean.

**Figure 4.**
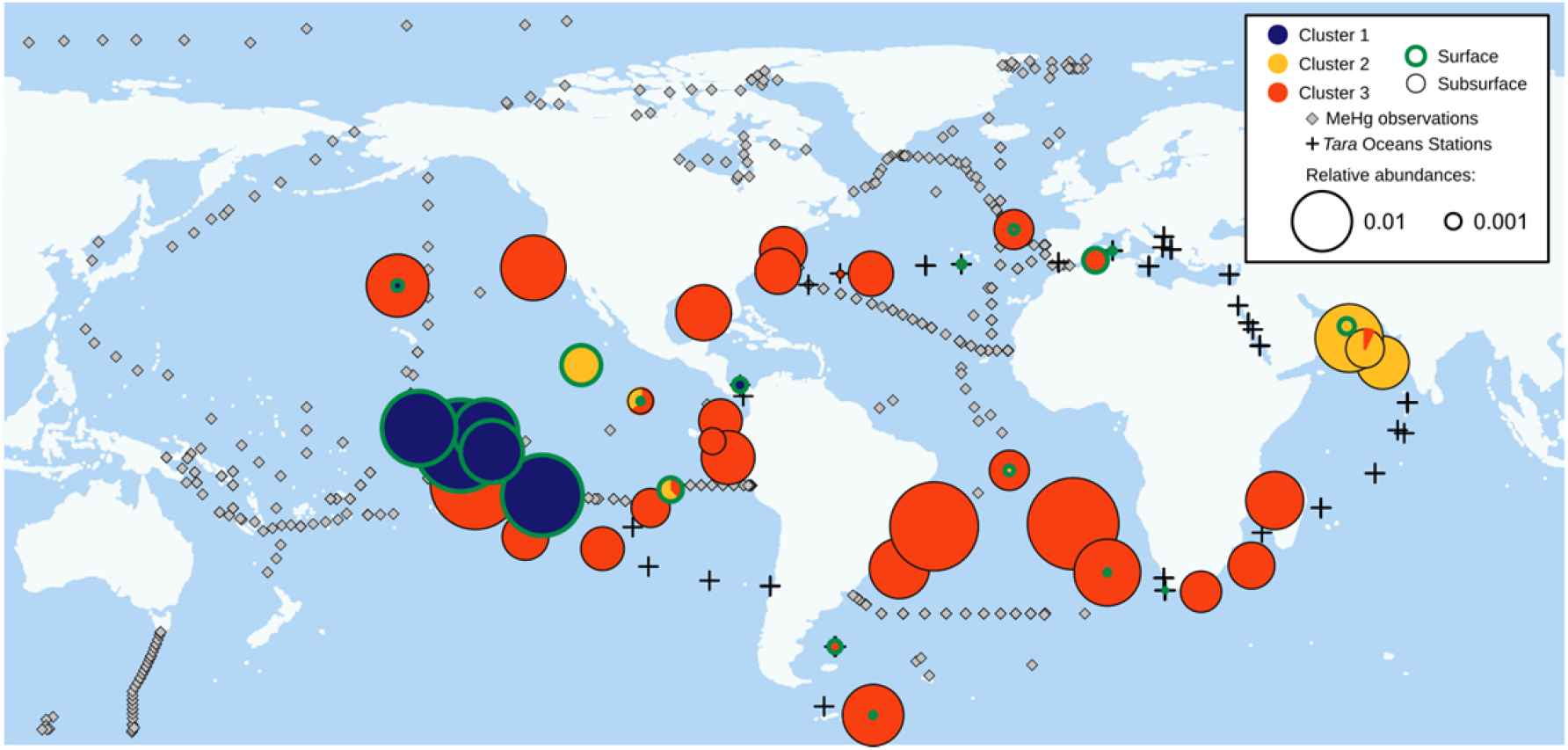
HgcA biogeography. Circle sizes are proportional to the cumulated HgcA homolog genes abundances at each station. The pie charts indicate the cluster attribution (legend in the chart), and their border color indicates the sampling depth: surface samples (<120 m-depth) in green and subsurface samples (> 120 m-depth) in grey. *Tara* Oceans stations without detected *HgcAB* genes are represented by black crosses and seawater MeHg profiles from the literature (Supplementary Text 1) by black diamonds.

### Nitrospina as the most predominant and widespread methylator in the open ocean

The predominant and widespread HgcA-like homologs were phylogenetically extremely close to the *Nitrospina*-related ones (Supplementary Figure 1) previously identified by metagenomic analysis as potential Hg-methylators within Antarctic sea ice and brine, and further detected by PCR in seawater samples below the ice ^13^. The four *Nitrospina* HgcA-like sequences from our study were distinct from HgcA in confirmed Hg-methylators, and also from HgcAB fusion proteins reported in environmental metagenomes ^12^ (Figure 2). The few cultured strains harboring a fused *hgcAB* gene (*Methanococcoides methylutens* and *Pyrococcus furiosus)* were unable to produce MeHg in experimental conditions ^12,27^. Through sequence alignment against Protein Data Bank templates, we confirmed that the four *Nitrospina* HgcA-like homologs showed high conservation of six residue positions involved in cobalamin binding, which is mandatory for methyl group transfer to Hg ^13^ (Supplementary Fig. 1). Protein structure modelling suggests that some *Nitrospina* species may be capable of Hg-methylation. The observed mutations (N71 and C74) do not suppress Hg methylation capacity, according to mutagenesis experiments in the model methylator *D. desulfuricans* ND132 ^15^. The strictly conserved cysteine facilitates the transfer of methyl groups to inorganic Hg ^28^. The two current *Nitrospina* isolates (*N. gracilis* and *N. watsonii*) have not been experimentally tested to date for their Hg-methylation capacity. *N. gracilis* genome lacks the *hgcA* gene. The complete genome of *N. watsonii* is not available. From the 12 *Nitrospina* genome assemblies available on NCBI at the time of writing, we found HgcA-like proteins (harbouring the six mandatory residues for Hg-methylation) in three strains only: SCGC AAA288-L16 (single cell whole genome from 770 m-deep ALOHA station, North Pacific Ocean), AB-629-B06 (single cell whole genome from dark ocean water column) and LS_NOB (from a deep-sea sponge holobiont; Supplementary Fig. 1).

Mercury methylation has long been described for anaerobic environments ^12^ and *hgcA* genes have been found exclusively in anaerobic Bacteria and Archea ^5^. Yet, we find the most abundant HgcA homologs are strongly dominant in oxic subsurface samples, where they coincide with the subsurface MeHg concentration peaks ^6^, and are carried by the nitrite oxidizing bacteria *Nitrospina*, usually considered as aerobic.

Several clues may explain this apparent contradiction. First, it is increasingly recognized that anaerobic processes can occur in anoxic niches such as organic matter aggregates in the middle of oxic waters ^8^. *Nitrospina* sequences were predominantly present in the larger size fractions (accounting for 78% of total HgcA abundances), suggesting that Hg-methylation -as other anaerobic processes-might be associated with particles, where anoxic niches are likely to be favourable to *Nitrospina* methylating activity. Several features suggest the adaptation of *Nitrospina* to low-oxygen environments. *Nitrospina* have been detected as particularly abundant (up to 10% of the bacterial community) in several upwelling and oxygen-deficient zones ^29^.

Comparative genomics revealed a close evolutionary relationship between *Nitrospina* and Anammox bacteria, including horizontal gene transfer events, suggesting the coexistence of these organisms in hypoxic or anoxic environments ^16^, as confirmed in incubation experiments ^30,31^. Genome analysis of several *Nitrospina* strains revealed unexpected adaptation features to low-oxygen environments: no ROS defence mechanism, dependence on highly oxygen-sensitive enzymes for carbon fixation, and high O_2_-affinity cytochromes ^16,32^.

*Nitrospina* can play diverse ecological roles beyond the nitrogen cycle ^33^. *Nitrospina* can use alternative anaerobic pathways to gain energy, using other terminal electron acceptors than O_2_ during fermentation under hypoxic or anaerobic conditions, such as sulfur compounds or metal oxides. Their capacity to cope with environmental Hg through methylation is worth considering, since their genome is well equipped against other toxic compounds (arsenate- and mercuric-reductase, metallic cation transporters, multidrug export system) ^16,32^. Mercury methylation potential might have been acquired by horizontal gene transfer. Within the four *Nitrospina* scaftigs harbouring *hgcA*, other neighbour genes related to methyl group transfer and Hg metabolism are found, such as the *merR1* regulator of the *mer* operon involved in Hg resistance, the *ubiE* methyltransferase and the putative metal-binding YHS domain (Fig. 1). This genomic context can lead to hypothesize that the expression of these genes, including *hgcA*, is under the same Hg-induced regulation as the *mer* operon, triggered by *merR*.

The choice of *hgcAB* as an indicator of Hg-methylation has to be discussed. First, the presence of *hgcAB* appears necessary but not sufficient for Hg methylation. Indeed, unsuccessful attempts to transfer Hg-methylation capacity to a non-Hg-methylating strain suggest that unidentified additional genes might be needed for effective MeHg production 15. Several critical steps are involved in the Hg-methylation process, such as Hg(II) sensing, cellular uptake of Hg(II) by active transport, methyl-group providing and transfer, and MeHg export from the cell. All these steps could be targeted as functional markers of Hg-methylation in the environment in order to provide a more complete picture of the process. Second, the exact contribution of HgcAB to Hg-methylation is not well understood. Since Hg methylation does not confer Hg resistance, it cannot be considered as a protection mechanism against Hg toxicity ^19^. In model strains *D. dechloroacetivorans*, net Hg methylation was not clearly induced by inorganic Hg and not significantly correlated to *hgcAB* gene expression levels, but rather influenced by environmental factors, growth conditions and energetic metabolism ^19,34^. The variability of the methylation potential has been evidenced in different strains, and the implication of *hgcAB* might also vary between strains. Such functional gene approaches are powerful to track biogeochemical potentials in extended environments but remain limited to well described metabolic pathways, ignoring genes with unknown function ^35^.

Here, we bring metagenomic evidence for widespread presence of microbial Hg-methylators in the global ocean, thus reconciling with previous geochemical hints pointing to *in situ* MeHg production in the water column. The key Hg-methylating genes found across all oceans corresponded to taxonomic relatives of known Hg-methylators from Deltaproteobacteria, Firmicutes and Chloroflexi phyla. We further identified the microaerophilic NOB *Nitrospina* as the potential dominant Hg-methylator in the global ocean, ubiquitous at the DNA-level, and favoured by oxic subsurface waters (Supplementary Figure 2). A critical next step would be to examine their *hgcA* expression levels and to evaluate Hg-methylation capacity in *Nitrospina* cultures. Further studies should also determine the physicochemical parameters controlling *Nitropina* Hg-methylation activity level, in order to better understand how they will respond to expected global changes. Our results open new avenues for disentangling the functional role of microorganisms in marine Hg cycling. Our analysis of the *Tara* Oceans metagenomes reveals global distribution of the key Hg methylating genes (*hgcA* and *hgcB*) and pinpoints *Nitrospina* as responsible for widespread open ocean MeHg production in subsurface oxic seawater. Our study implicates the subsurface oxic waters of all oceans as potential source of MeHg that should be considered in the global Hg-cycle budgets, and identifies microbial target for further research on marine MeHg production. We hypothesize that besides anthropogenic Hg emissions, ongoing global climate change might have a previously underestimated effect on *in situ* marine MeHg production by water-column microorganisms, by disturbing microbial assemblages, activity, and environmental drivers governing Hg-methylation.

## Methods

### Identification of HgcAB environmental sequences in oceanic metagenomes

*hgcA* and *hgcB* genes encode for a putative corrinoid protein, HgcA, and a 2[4Fe-4S] ferredoxin, HgcB, serving respectively as methyl group carrier and electron donor for corrinoid cofactor reduction. HgcA and HgcB homologs were retrieved by searching Hidden Markov Model profiles (HMM) ^36^ provided by Podar et al. ^12^ in the Ocean Microbial Reference Gene Catalog^14^ (OM-RGC) using the Ocean Gene Atlas^37^ (http://tara-oceans.mio.osupytheas.fr/ocean-gene-atlas/). The OM-RGC is the most exhaustive catalogue of marine genes to date including datasets from *Tara* Oceans metagenomic assemblies and other publicly available marine genomic and metagenomic datasets. We applied an e-value threshold of 1e-20. The corresponding scaftigs (i.e. the assembled sequences where the homolog genes were predicted) were retrieved from the raw assemblies deposited at ENA (Supp Data 1 & 4). Eight scaftigs without *Tara* Oceans mapped reads were discarded. The remaining scaftigs were annotated using Prokka with default parameters ^38^. The resulting translated sequences were aligned separately for HgcA and HgcB using Jalview 2.10 and alignments were cleaned manually^39^. For further analysis, we kept HgcA sequences if they possess the conserved motif NVWCAA^5^, or if the neighbouring HgcB sequence was present on the scaftig.

### HgcA phylogenetic analysis

A phylogenetic tree was built from the 10 HgcA sequences kept, 55 HgcA protein sequences representative of known Hg-methylator clades belonging to Archaea, Firmicutes, Chloroflexi and Deltaproteobacteria, including 18 experimentally-confirmed Hg-methylators (initially published by Parks et al. ^5^, and updated at https://www.esd.ornl.gov/programs/rsfa/data.shtml), as well as 9 HgcAB fusion proteins ^13^ and 3 HgcA-like proteins predicted from *Nitrospina* genome assemblies using Prokka ^38^. The tree was rooted with 3 paralogs from confirmed non-Hg-methylating strains ^13^. The closest sequences (i.e. best e-value match) of each environmental HgcA sequence were retrieved using BLASTp against non-redundant RefSeq protein database excluding sequences from uncultured organisms ^40^, and included in the tree.

The 80 sequences were aligned using MAFFT ^41^ and gap-containing sites were removed using the mode gappyout of TrimAl ^42^. Maximum likelihood phylogenies were inferred using PhyML Best AIC Tree (version 1.02b) implemented in Phylemon^43^ (version 2.0) with the best model of sequence evolution Blosum62+I+G+F. Branch support was calculated using the non-parametric Shimodaira-Hasegawa-like approximate likelihood ratio test. The final tree was edited using Evolview ^44^, especially by annotating the isolation sites retrieved from Genomes OnLine Database ^45^.

### Conserved sites in HgcA

Four sequences from OM-RGC related to *Nitrospina* were aligned with the same Protein Data Bank (PDB) templates as Gionfriddo et al. ^13^, as well as the 3 HgcA-like proteins from *Nitrospina* genome assemblies, and conserved residues were checked. The chosen PDB structural templates (4djd_C, 2h9a_A, 4C1n_C, 2ycl_A) were the gamma subunit of the corrinoid S-Fe acetyl-CoA decarbonylase/synthase complex, identified by Gionfriddo et al. ^13^ as the closest and most complete relative to currently unresolved HgcA structure.

### Biogeography of HgcA

Relative HgcA abundances in *Tara* Oceans samples were obtained from the Ocean Gene Atlas^37^. We screened 243 metagenomes from 68 sites covering the World Ocean except Arctic, sampled at different depths from surface to 500 m-depth, covering six different size fractions ranging from 0 to 3 µm. Environmental data were obtained from Pesant et al. ^46^ (Supplementary Table 2). For the following analysis, we considered two depths classes (surface samples < 120 m-depth, subsurface samples > 120 m-depth), two particle size fractions (< 5 µm, < 0.8 µm), two oxic states (oxic: O_2_ > 10 µM, suboxic: O_2_ < 10 µM). HgcA relative abundance was calculated as follows: the length-normalized count of genes read was divided by the median of the length-normalized counts of a set of ten universal single copy marker genes ^47,48^. Thus, relative abundance represents the fraction of bacteria harbouring the *hgcA* gene within the assembled genomes. A heatmap of the relative gene abundances in *Tara* Oceans samples was generated in R ^49^ using the heatmap.2 function in the ggplot CRAN library. Dendrograms were computed using hclust default parameters from Ward distance index based on presence/absence of the genes (‘binary’ option). Genes were clustered into three groups (Cluster 1, Cluster 2 and Cluster 3) according to their abundance pattern on the heatmap. The geographic origin of the *hgcA* genes retrieved from the *Tara* Oceans samples was plotted on a global map using the “mapplots” R package. At each station, the cumulated abundance and phylogenetic affiliation of the retrieved *hgcA* genes were represented on the map by the size and colour of the points. Cluster distribution was also plotted against depth and oxygen concentration at each station to depict the environmental conditions where each Cluster flourishes (Supplementary Figure 2). Tracks of MeHg records from previous campaigns were searched in the literature (Supplementary Text 1) and georeferenced on the map.

## Acknowledgements

The authors thank Patricia Bonin, Joana R.H. Boavida, Pascal Hingamp, Eric Pelletier, Daniel Cossa, Jeroen E. Sonke for constructive comments that helped to improve this manuscript.

## Author contributions

E.V., L.C. and L.E.H.B. wrote the manuscript. E.V. performed the bioinformatic analyses with the scientific support of L.C.

## References

1. Outridge, P. M., Mason, R. P., Wang, F., Guerrero, S. & Heimbürger-Boavida, L. E.Updated Global and Oceanic Mercury Budgets for the United Nations Global Mercury Assessment 2018. Environ. Sci. Technol. 52, 11466–11477 (2018).

2. Gilmour, C. C. et al. Mercury methylation by novel microorganisms from new environments. Environ. Sci. Technol. 47, 11810–11820 (2013).

3. Gilmour, C. C. et al. Sulfate-Reducing Bacterium Desulfovibrio desulfuricans ND132 as a Model for Understanding Bacterial Mercury Methylation. Appl. Environ. Microbiol. 77, 3938–3951 (2011).

4. Mason, R. P. et al. Mercury biogeochemical cycling in the ocean and policy implications. Environ. Res. 119, 101–117 (2012).

5. Parks, J. M. et al. The genetic basis for bacterial mercury methylation. Science (80-.). 339, 1332–1335 (2013).

6. Schlitzer, R. et al. The GEOTRACES Intermediate Data Product 2017. Chem. Geol. 493, 210–223 (2018).

7. Ortiz, V. L., Mason, R. P. & Evan Ward, J. An examination of the factors influencing mercury and methylmercury particulate distributions, methylation and demethylation rates in laboratory-generated marine snow. Mar. Chem. 177, 753–762 (2015).

8. Bianchi, D., Weber, T. S., Kiko, R. & Deutsch, C.Global niche of marine anaerobic metabolisms expanded by particle microenvironments. Nat. Geosci.1–6 (2018). doi:10.1038/s41561-018-0081-0

9. Lehnherr, I., St. Louis, V. L., Hintelmann, H. & Kirk, J. L.Methylation of inorganic mercury in polar marine waters. Nat. Geosci. 4, 298–302 (2011).

10. Blum, J. D.Marine mercury breakdown. Nat. Geosci. 4, 139–140 (2011).

11. Masbou, J. et al. Carbon Stable Isotope Analysis of Methylmercury Toxin in Biological Materials by Gas Chromatography Isotope Ratio Mass Spectrometry. Anal. Chem. 87, 11732–11738 (2015).

12. Podar, M. et al. Global prevalence and distribution of genes and microorganisms involved in mercury methylation. Sci. Adv. 1, (2015).

13. Gionfriddo, C. M. et al. Microbial mercury methylation in Antarctic sea ice. Nat. Microbiol. 1, 16127 (2016).

14. Sunagawa, S. et al. Ocean plankton. Structure and function of the global ocean microbiome. Science 348, 1261359 (2015).

15. Smith, S. D. et al. Site-directed mutagenesis of HgcA and HgcB reveals amino acid residues important for mercury methylation. Appl. Environ. Microbiol. 81, 3205–17 (2015).

16. Lücker, S., Nowka, B., Rattei, T., Spieck, E. & Daims, H.The Genome of Nitrospina gracilis Illuminates the Metabolism and Evolution of the Major Marine Nitrite Oxidizer. Front. Microbiol. 4, 27 (2013).

17. Spieck, E., Keuter, S., Wenzel, T., Bock, E. & Ludwig, W.Characterization of a new marine nitrite oxidizing bacterium, Nitrospina watsonii sp. nov., a member of the newly proposed phylum “Nitrospinae”. Syst. Appl. Microbiol. 37, 170–176 (2014).

18. Cao, J. et al. Pseudodesulfovibrio indicus gen. nov., sp. nov., a piezophilic sulfate-reducing bacterium from the Indian Ocean and reclassification of four species of the genus Desulfovibrio. Int. J. Syst. Evol. Microbiol. 66, 3904–3911 (2016).

19. Gilmour, C. C. et al. Sulfate-reducing bacterium Desulfovibrio desulfuricans ND132 as a model for understanding bacterial mercury methylation. Appl. Environ. Microbiol. 77, 3938–51 (2011).

20. Bae, H.-S., Dierberg, F. E. & Ogram, A.Syntrophs dominate sequences associated with the mercury methylation-related gene hgcA in the water conservation areas of the Florida Everglades. Appl. Environ. Microbiol. 80, 6517–26 (2014).

21. Sorokin, D. Y., Tourova, T. P., Mußmann, M. & Muyzer, G.Dethiobacter alkaliphilus gen. nov. sp. nov., and Desulfurivibrio alkaliphilus gen. nov. sp. nov.: two novel representatives of reductive sulfur cycle from soda lakes. Extremophiles 12, 431–439 (2008).

22. Tidjani Alou, M. et al. Gut Bacteria Missing in Severe Acute Malnutrition, Can We Identify Potential Probiotics by Culturomics? Front. Microbiol. 8, 899 (2017).

23. Regnell, O. & Watras, C. J.Microbial Mercury Methylation in Aquatic Environments: A Critical Review of Published Field and Laboratory Studies. Environ. Sci. Technol. 53, 4–19 (2019).

24. Sela-Adler, M. et al. Co-existence of Methanogenesis and Sulfate Reduction with Common Substrates in Sulfate-Rich Estuarine Sediments. Front. Microbiol. 8, 766 (2017).

25. Pak, K. R. & Bartha, R.Mercury methylation by interspecies hydrogen and acetate transfer between sulfidogens and methanogens. Appl. Environ. Microbiol. 64, 1987–1990 (1998).

26. Roullier, F. et al. Particle size distribution and estimated carbon flux across the Arabian Sea oxygen minimum zone. Biogeosciences 11, 4541–4557 (2014).

27. Gilmour, C. C., Bullock, A. L., McBurney, A., Podar, M. & Elias, D. A.Robust mercury methylation across diverse methanogenic Archaea. MBio 9, (2018).

28. Zhou, J., Riccardi, D., Beste, A., Smith, J. C. & Parks, J. M.Mercury Methylation by HgcA: Theory Supports Carbanion Transfer to Hg(II). Inorg. Chem. 53, 772–777 (2014).

29. Levipan, H. A., Molina, V. & Fernandez, C.Nitrospina-like bacteria are the main drivers of nitrite oxidation in the seasonal upwelling area of the Eastern South Pacific (Central Chile ∼36°S). Environ. Microbiol. Rep. 6, 565–573 (2014).

30. Füssel, J. et al. Nitrite oxidation in the Namibian oxygen minimum zone. ISME J. 6, 1200–1209 (2012).

31. Beman, J. M., Leilei Shih, J. & Popp, B. N.Nitrite oxidation in the upper water column and oxygen minimum zone of the eastern tropical North Pacific Ocean. ISME J. 7, 2192–2205 (2013).

32. Ngugi, D. K., Blom, J., Stepanauskas, R. & Stingl, U.Diversification and niche adaptations of Nitrospina-like bacteria in the polyextreme interfaces of Red Sea brines. ISME J. 10, 1383–1399 (2016).

33. Daims, H., Lücker, S. & Wagner, M.A New Perspective on Microbes Formerly Known as Nitrite-Oxidizing Bacteria. Trends Microbiol. 24, 699–712 (2016).

34. Goñi-Urriza, M. et al. Relationships between bacterial energetic metabolism, mercury methylation potential, and hgcA/hgcB gene expression in Desulfovibrio dechloroacetivorans BerOc1. Environ. Sci. Pollut. Res. 22, 13764–13771 (2015).

35. Reed, D. C., Algar, C. K., Huber, J. A. & Dick, G.J. Gene-centric approach to integrating environmental genomics and biogeochemical models. Proc. Natl. Acad. Sci. U. S. A. 111, 1879–84 (2014).

36. Eddy, S. R.Accelerated profile HMM searches. PLoS Comput. Biol. 7, e1002195 (2011).

37. Villar, E. et al. The Ocean Gene Atlas: exploring the biogeography of plankton genes online. Nucleic Acids Res. 46, W289–W295 (2018).

38. Seemann, T. Prokka: rapid prokaryotic genome annotation. Bioinformatics 30, 2068–2069 (2014).

39. Waterhouse, A. M., Procter, J. B., Martin, D. M. A., Clamp, M. & Barton, G. J.Jalview Version 2—a multiple sequence alignment editor and analysis workbench. Bioinformatics 25, 1189–1191 (2009).

40. Altschul, S. F., Gish, W., Miller, W., Myers, E. W. & Lipman, D. J.Basic local alignment search tool. J. Mol. Biol. 215, 403–410 (1990).

41. Katoh, K. & Standley, D. M.MAFFT multiple sequence alignment software version 7: improvements in performance and usability. Mol. Biol. Evol. 30, 772–780 (2013).

42. Capella-Gutierrez, S., Silla-Martinez, J. M. & Gabaldon, T.trimAl: a tool for automated alignment trimming in large-scale phylogenetic analyses. Bioinformatics 25, 1972–1973 (2009).

43. Sanchez, R. et al. Phylemon 2.0: a suite of web-tools for molecular evolution, phylogenetics, phylogenomics and hypotheses testing. Nucleic Acids Res. 39, W470–W474 (2011).

44. Gao, S. et al. Evolview v2: an online visualization and management tool for customized and annotated phylogenetic trees. Nucleic Acids Res. 44, W236–W241 (2016).

45. Mukherjee, S. et al. Genomes OnLine Database (GOLD) v.6: data updates and feature enhancements. Nucleic Acids Res. 45, D446–D456 (2017).

46. Pesant, S. et al. Open science resources for the discovery and analysis of Tara Oceans data. Sci. Data 2, 150023 (2015).

47. Mende, D. R., Sunagawa, S., Zeller, G. & Bork, P.Accurate and universal delineation of prokaryotic species. Nat. Methods 10, 881–884 (2013).

48. Sunagawa, S. et al. Metagenomic species profiling using universal phylogenetic marker genes. Nat. Methods 10, 1196–1199 (2013).

49. R Development Core Team. R: A Language and Environment for Statistical Computing.(2008).

